# Remorin proteins serves as membrane topology scaffolds in plants

**DOI:** 10.1101/2022.03.15.484384

**Authors:** Chao Su, Marta Rodriguez-Franco, Beatrice Lace, Nils Nebel, Casandra Hernandez-Reyes, Pengbo Liang, Eija Schulze, Claudia Popp, Jean Keller, Cyril Libourel, Alexandra A.M. Fischer, Katharina E. Gabor, Evgeny V. Mymrikov, Nikolas M. Gross, Eric Mark, Carola Hunte, Wilfried Weber, Petra Wendler, Thomas Stanislas, Pierre-Marc Delaux, Oliver Einsle, Thomas Ott

## Abstract

Organization of membrane topologies in plants has so far been mainly attributed to the cell wall and the cytoskeleton. Taking rhizobial infections of legume root cells, where plasma membranes undergo dynamic and large-scale topology changes, as an initial model, we challenged this paradigm and tested whether additional scaffolds such as plant-specific remorins that accumulate on highly curved and often wall-less plasma membrane domains, control local membrane dynamics. Indeed, loss-of-function mutants of the remorin protein SYMREM1 failed to develop stabilized membrane tubes as found in colonized cells in wild-type plants, but released empty membrane spheres instead. Expression of this and other remorins in wall-less protoplasts allowed engineering different membrane topologies ranging from membrane blebs to long membrane tubes. Reciprocally, mechanically induced membrane indentations were equally stabilized by SYMREM1. This function is likely supported by remorin oligomerization into antiparallel dimers and the formation of higher order membrane scaffolding structures. Taken together we describe an evolutionary confined mechanism that allows the stabilization of large-scale membrane conformations and curvatures in plants.

**One-sentence summary:** The remorin SYMREM1 evolved as structural membrane scaffold that stabilizes membrane tubulation and curvature during symbiotic intracellular infections.

## Introduction

Intracellular colonization of host cells is a central feature of mutualistic associations like the root nodule symbiosis (RNS) and the arbuscular mycorrhiza symbiosis that occur between a host plant and soil-borne rhizobia or Glomeromycotean fungi, respectively. Molecularly, the RNS-related infection is tightly controlled and requires the perception of strain-specific rhizobial lipochitooligosaccharides (Nod factors) by host LysM-type receptor-like kinases ^1-3^ and of rhizobial exopolysaccharides ^4^. Perception of these microbial patterns by heteromeric receptor complexes trigger a symbiotic signaling cascade that results in perinuclear calcium spiking ^5^. This calcium signature is, in turn, decoded by the calcium-calmodulin-dependent kinase CCaMK/DMI3 and the transcriptional activator CYCLOPS/IPD3 ^6,7^. Upon phosphorylation of CYCLOPS/IPD3 by CCaMK/DMI3, RNS-specific gene expression is triggered by the activation of specific transcription factors such as NODULE INCEPTION (NIN) ^8,9^. Consequently, mutations in these genes results in the inability of the host to maintain intracellular infections and bacterial release.

Intruding rhizobia mostly infect legume hosts such as *Medicago truncatula* and *Lotus japonicus* via young, growing root hairs that swell and later curl around surface attached rhizobia to entrap them just below the root hair tip. This process is driven by cellular repolarization of the actin and microtubule cytoskeleton ^10-13^ and involves, among others, the actin polymerizing formin protein SYFO1 ^14^. Root hair curling results in an entrapment of the symbiont in a so-called infection chamber ^15^. A re-polarization of the cell towards the infection chamber leads to targeted secretion of proteins and membrane constituents, which enables the formation of a membrane-surrounded tunnel called the ‘infection thread’ (IT) that emerges from the infection chamber, then transcellularly progresses though the root cortex and finally branches inside the nodule primordium ^16,17^. Within these primordia and later in indeterminate nodules of *M. truncatula*, rhizobia are continuously released from bulges of nodular ITs (infection droplets) into these infected cells. Upon release, rhizobia differentiate into nitrogen-fixing bacteroids that remain encapsulated by the host-derived symbiosome/peribacteroid membrane ^18,19^. During infection, initial membrane invaginations, young IT segments around the growing tip, infection droplets as precursors of bacterial release sites, and symbiosome membranes encapsulating the nitrogen-fixing symbiont are devoid of a rigid cell wall that could provide structural support to these sites ^20,21^.

In many organisms, membrane topologies can be maintained by oligomeric scaffold proteins like clathrins as described during endocytosis ^22,23^ or Bar-domain proteins like amphisin or BIN1 in human cells ^24^. However, our mechanistic understanding of membrane stabilization in the absence of a primary cell wall in plant cells is rather sparse.

During RNS, the scaffold SYMREM1, a member of the plant-specific remorin family ^25^, recruits and stabilizes the symbiotic receptor LYK3 in membrane nanodomains ^26^. This function might be explained by remorin-induced alterations in membrane fluidity ^27,28^, higher order protein oligomerization ^29-31^ or maintenance of membrane-associated and phase-separated condensates ^32^. Like all other remorins, SYMREM1 is comprised of a conserved alpha-helical C-terminal region, while the intrinsically disordered N-terminal region (IDR) of SYMREM1 is highly variable in sequence and conformation ^33^. Membrane association of remorins is mediated by the Remorin C-terminal Anchor (RemCA) and, in most cases, assisted by palmitoylation ^33,34^. Furthermore, it has been shown that remorins oligomerize at the plasma membrane *in planta* ^35^ and can form higher order filamentous structures or protein lattices *in vitro* ^31^.

Since remorins can alter membrane fluidity ^28^ and *symrem1* mutants largely fail to release rhizobia into nodule cells ^36^, these proteins may have greater impact in membrane topology than currently envisioned. Furthermore, SYMREM1 accumulates predominantly on cell wall-devoid symbiotic membranes such as IT tips ^26^, nodular infection droplets that precede bacterial release and the symbiosome membrane, which surrounds the released and nitrogen-fixing rhizobia inside the nodule ^36^. Therefore, we investigated putative roles of SYMREM1 in membrane dynamics and shape.

## Results

To visualize symbiotic membranes in great detail and to examine their precise morphology, we labelled phosphatidylserine (PS), a central phospholipid of biological membranes, by expressing a LactC2 biosensor ^37^. This allowed clear imaging of membrane patterns at ITs, infection droplets and symbiosome membranes (Fig. S1). While ITs and infection droplet structures were mostly filled with rhizobia, we frequently observed additional empty tube-like membrane structures on enlarged ITs associated with infection droplets (Fig. 1A) in nodule cortex cells of wild-type plants. As previously shown and recapitulated here, all three membrane sites mentioned above were also targeted by SYMREM1 (Fig. 2A-2G) ^36^. In line with this accumulation of SYMREM1 at ITs, on infection droplets and on symbiosome membranes, and in contrast to WT plants, *symrem1* mutants failed to release bacteria in a majority of cells in the inner nodule cortex and exhibited bulky ITs, as revealed by light and electron microscopy (Fig. 2H-2M) ^36^. To assess possible alternations in membrane topology, we also expressed the PS-biosensor in the *symrem1-2* mutant. In contrast to WT plants, ITs observed in the *symrem1* mutant did not display signs of membrane tubulation (Fig. 1B). Instead, large amounts of detached empty membrane spheres were observed in IT-containing but release-deficient nodule cortex cells of *symrem1* mutants as assessed by confocal laser-scanning microscopy (Fig. 1C and 1D) and transmission electron microscopy (TEM) (Fig. 1F-1H). This is in sharp contrast to WT plants, where bacterial differentiation and symbiosome formation occurred normally with symbiosome membranes being tightly aligned with the differentiated bacteroids (Fig. 1E, Fig. S1F). These data together with the fact that the entry receptor LYK3 does not localizes to the infection droplet membrane ^38^ implied that SYMREM1 may not only serve as a scaffold for symbiotic receptors but possibly serves additional roles in maintaining a functional topology of symbiotic membranes, which are not delimited by a modified cell wall. The hypothesis of SYMREM1 being a structural membrane scaffold was further supported by the fact that other members of the remorin family have been shown to form higher order proteinaceous lattices and/or filamentous structures ^30,31,39,40^ *in vitro*. We also confirmed such filamentous structures for purified, recombinant SYMREM1 using TEM analysis followed by 2D classification of manually picked filamentous segments. Here, we detected auto-assembled amorphous protein filaments that were partially branched or scrambled (Fig. 3A). Systematic inspection of 389 filament fragments revealed an average width between 84 and 125 Å for the filamentous particles with a few class averages showing helical features such as twists (Fig. 3B). Besides this, we mostly observed irregular filaments and protein bodies (Fig. 3A) that may represent filament seeds. Analytical gel-filtration of fresh SYMREM1 protein extracts revealed an apparent molecular weight of 57 kDa that matches the calculated size for a dimeric remorin protein (Fig. 3C). Since the filaments were too amorphous for further structural assessment by cryo-EM, we conducted 3D modelling using Alphafold ^41^. In line with previous analyses, the algorithm predicted a largely disordered N-terminal region encompassing residues 1-69, followed by an α-helical segment from residues 70-187 and a disordered C-terminus from 188-205. Interestingly, the search for multimers of SYMREM1 consistently yielded a head-to-tail dimer as a recurrent structural unit (Fig. 3D). Here, the slightly bent α-helical segments of two monomers interacted through extended hydrophobic patches, with the N-termini pointing outward. Using Alphafold2 ^42^ predictions for higher order oligomers we repeatedly obtained alignments in flexible sheets that can be extended to helical structures (Fig. 3E). Our model further indicated a consistent interaction between two dimers that was not mediated by hydrophobic interactions, but rather by a few selected hydrogen bonds. Calculating electrostatic surface potentials on these structures revealed a strong enrichment of positive potentials at one site of the sheet, which further explains the tight association of SYMREM1 with the negatively charged head groups of plasma membrane lipids (Fig. 3F).

**Fig. 1.**
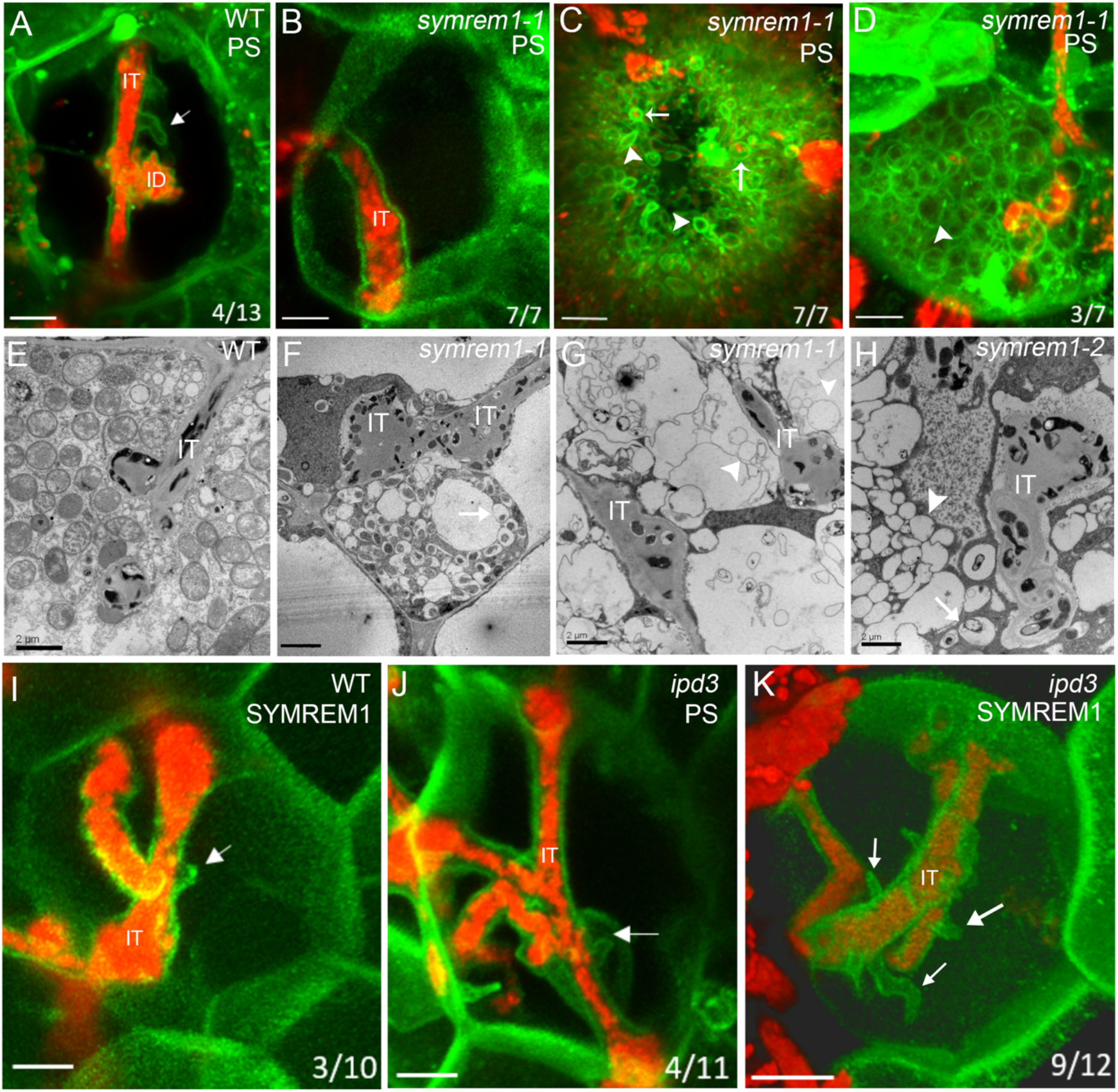
SYMREM1 stabilizes confined membrane tubes during infection. Spatially confined membrane tubes were found on wild-type (WT) IT (arrow, A) containing rhizobia (red) but not on ITs in *symrem1* mutants (B). Symbiosome membranes are loosely associated with released rhizobia (C) or appear as empty spheres (D) in *symrem1* mutants. Membranes were visualized by expressing the phosphatidylserine (PS) biosensor LactC2 (A-D). These patterns were confirmed by transmission electron microscopy for WT (E) and *symrem1* mutants (F-H). Arrows indicate symbiosome membranes that are loosely associated with released rhizobia and arrow heads indicate empty membrane spheres (C-H). Membrane tubes were found on nodular ITs in the *ipd3* mutant (J), while ectopic expression of SYMREM1 greatly increased these tubular outgrowths in the *ipd3* mutant (K) but not on WT (I). Arrows indicate SYMREM1-dependent membrane tubes in (I-K). Membranes were visualized by expressing the phosphatidylserine (PS) biosensor LactC2 (J) or YFP-SYMREM1 (I, K). Scale bars indicate 5 μm (A-D; I-K) and 2 μm (E-H). IT: infection thread; ID: infection droplet.

**Fig. 2:**
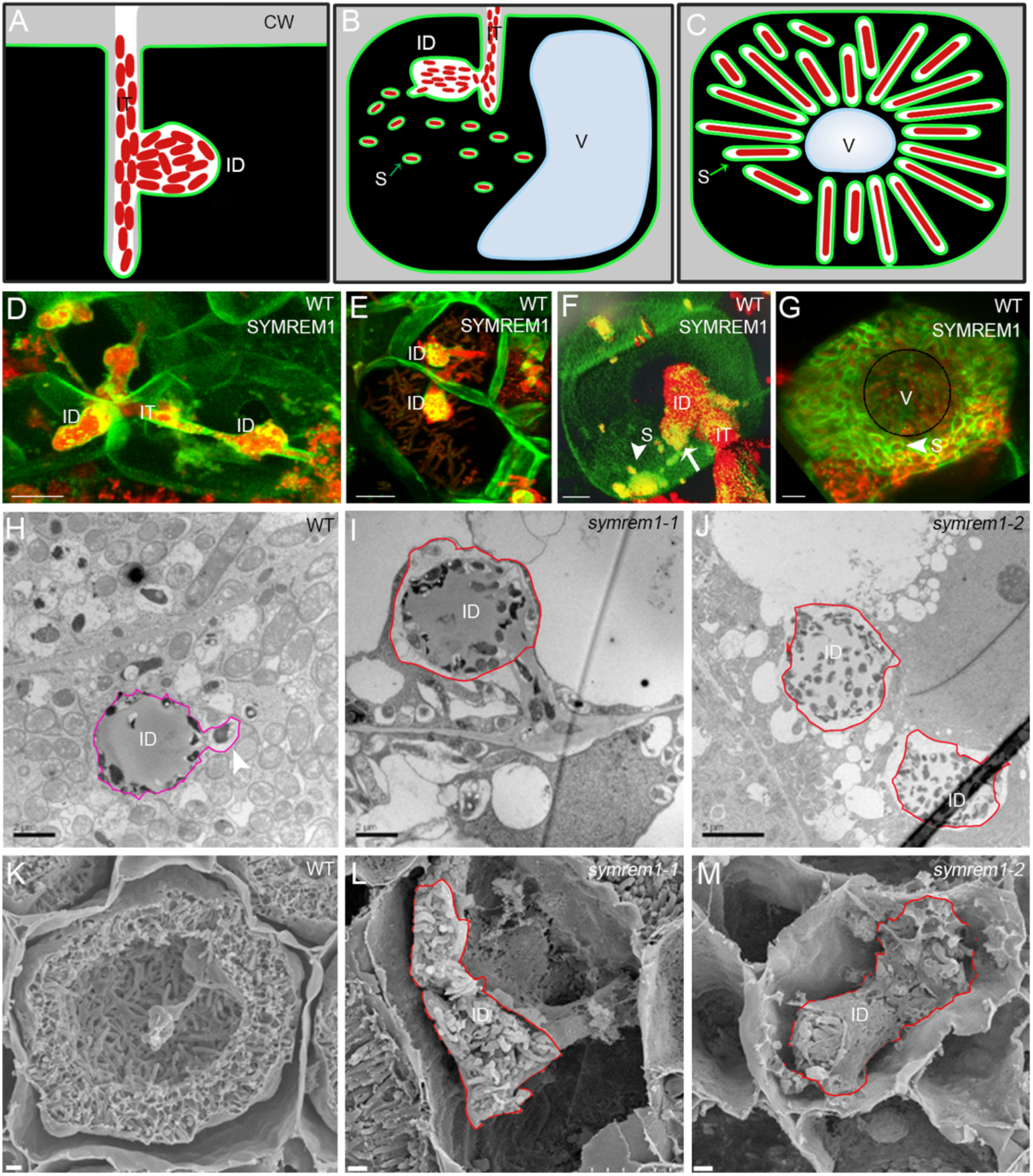
SYMREM1 functions in bacterial release from nodular infection threads. (A-C) Illustrations indicate an infection thread (IT) with an infection droplet (ID; A), and symbiosome (S) formation (B) and symbiosome-filled cell (C); CW: cell wall; V: vacuole. (D-G) Expression of an mEosEM-SYMREM1 fusion protein (green) specifically labels IT membranes, accumulates at bacterial droplet structures, and symbiosome membranes (*S. meliloti* expressing an mCherry marker; red). White arrow (in F) indicates the bacterial release site. (H-J) Transmission Electron Microscopy showing normal rhizobial release into wild-type (WT, H) while bacteria are trapped inside the IT droplet in *symrem1* mutants (I-J). ID, infection droplet (encircled). Scanning Electron Microscopy showing normal rhizobial release into bacteroids (K) while bacteria are trapped inside the infection droplet in *symrem1* mutants (L-M), Scale bars indicate 5 μm in (D-G and J), 2 μm in (H and I), 4 μm in K and 3 μm in L and M. Collapsed infection droplets in *symrem1* mutants were encircled with a red line (I, J, L and M).

**Fig. 3.**
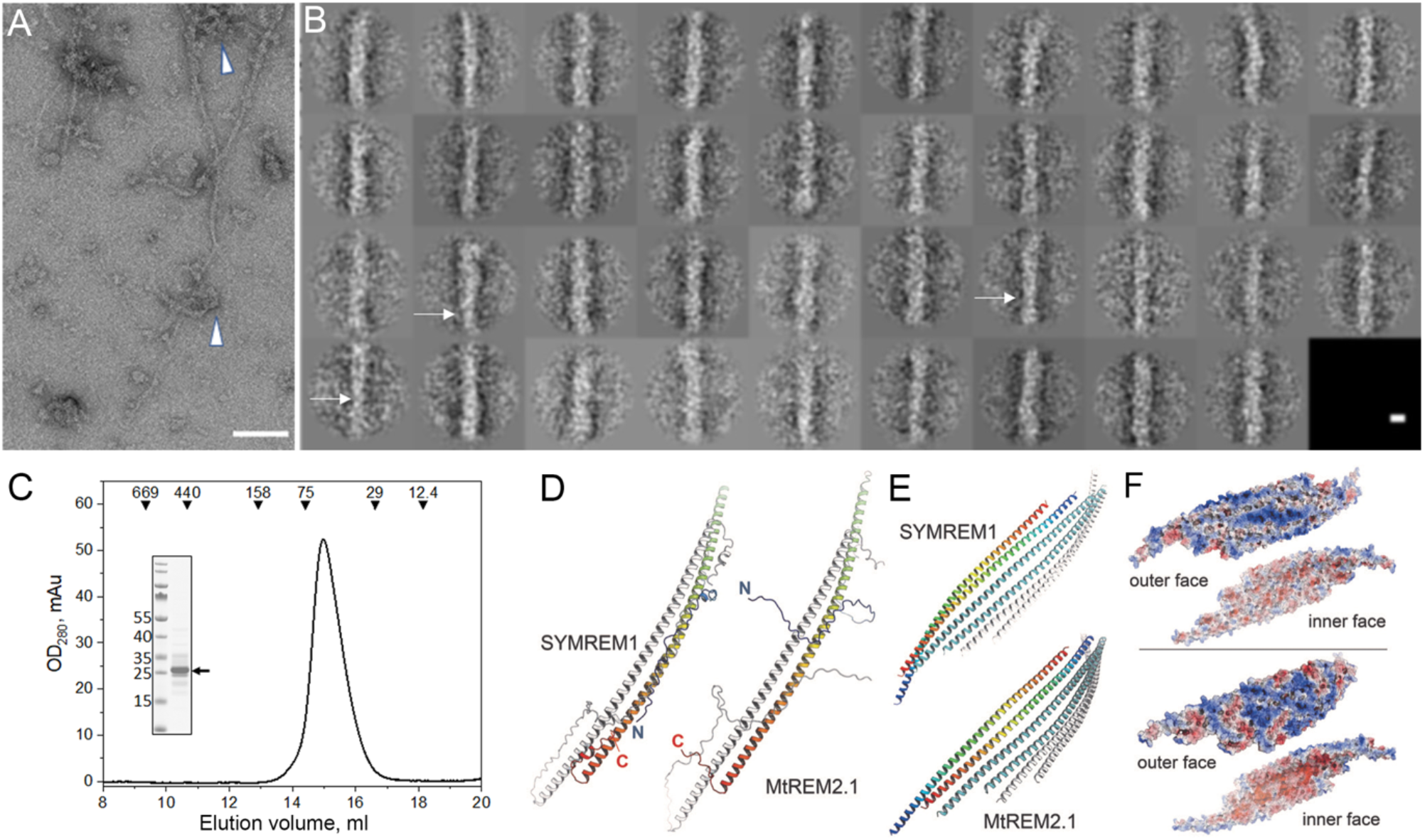
SYMREM1 forms oligomeric assemblies. (A) Representative raw electron micrograph of purified, recombinant SYMREM1 stained with 2% uranyl acetate. Arrow heads indicate irregular protein bodies. Scale bar indicates 100 nm. (B) 2D class averages derived from multivariate statistical analysis of all 389 particle images. Each class contains on average ten images. Class averages that show twisted features are marked with a white arrow. Scale bar indicates 100Å. (C) Elution profile of recombinant His-SYMREM1 purified on a Superdex200 Increase GL 10/30 column. Molecular masses (in kDa) and positions of elution peaks for standard proteins are indicated with triangles on the top. The elution volume of His-SYMREM1 was 15.0 ml, which corresponded to the apparent molecular mass of 56.9 kDa. Insert: SDS-PAGE after Coomassie staining of the purified His-SYMREM1 (labelled by an arrow); molecular masses of the pre-stained protein standards are indicated on the left in kDa. (D) In-silico predictions for homodimers of SYMREM1 (left) and MtREM2.1 (right). One monomer is colored from blue at the N-terminus to red at the C-terminus, the other in white. The Extended helical regions of both remorins form highly similar, antiparallel dimers. (E) Prediction of higher-order oligomers with Alphafold2. The remorin homodimers form flexible sheets that can be extended into helical structures. (F) Electrostatic surface potential maps for the two faces of the sheets formed by SYMREM1 (above) and MtREM2.1 (below), contoured from −5k_B_T (red) to +5k_B_T (blue). In both cases, the convex faces show a positive electrostatic potential, while that of the concave faces is negative.

The predicted sheet-like organization of the SYMREM1 oligomer with its positively charged surface prompted us to further test whether SYMREM1 has an impact on topological membrane features along infection threads. For this, we ectopically expressed fluorescently-labelled SYMREM1 in WT *M. truncatula* plants (Fig. 1I), a setup that was previously shown to elevate nodulation levels in *Lotus japonicus* ^43^. Here, we observed that the frequency of stabilized membrane tubes remained unaltered compared to those observed in LactC2-labelled WT cells (Fig. 1A and 1I), indicating a temporary nature of these structures at membrane interfaces that continuously release rhizobia. To test this hypothesis, we made use of the release-compromised *ipd3-1* mutant, which is defective in the transcriptional activator CYCLOPS that regulates, among other genes, the expression of endogenous *SYMREM1* ^44^. This reported transcriptional regulation also translated into reduced SYMREM1 protein levels in the *ipd3-1* mutant (Fig. S2). In line with this and similar to WT plants (Fig. 1A), PS-labelling revealed membrane tube formation in 36% of all assessed ITs in the *ipd3-1* mutant (Fig. 1J). However, ectopic expression of fluorophore-tagged SYMREM1 in *ipd3-1* significantly increased IT-associated membrane tubulation to 75% with several tubes per IT being frequently observed (Fig. 1K). These data further support a function of SYMREM1 in tubulation of membranes that are not supported by a rigid cell wall.

We then tested whether we can reconstitute SYMREM1-mediated effects on membrane topologies in cell wall-free *Nicotiana benthamiana* mesophyll protoplasts that adopt a spherical shape under mild hypotonic conditions as visualized by expressing a generic membrane marker probe (LTI6b-GFP; Fig S3A). However, when expressing SYMREM1, we observed numerous tubular outgrowths developing on 64 out of 112 (57%) inspected protoplasts (Fig. 4A-C, Fig S3B). Scanning electron microscopy revealed an average width of these protrusions of 178± 0.03 nm. This phenomenon was not observed on control protoplasts expressing the soluble and intrinsically disordered N-terminal region (IDR, amino acids 1-73) of the SYMREM1 protein (SYMREM1^IDR^; Fig. 4D, Fig. S3C and 3C’), while an N-terminally truncated SYMREM1 variant (SYMREM1^Cterm^, amino acids 74-205) that lacked the IDR but supported the higher order oligomer in our AlphaFold predictions, maintained the ability to stabilize membrane tubes at the protoplast surface (Fig. 4E).

**Fig. 4:**
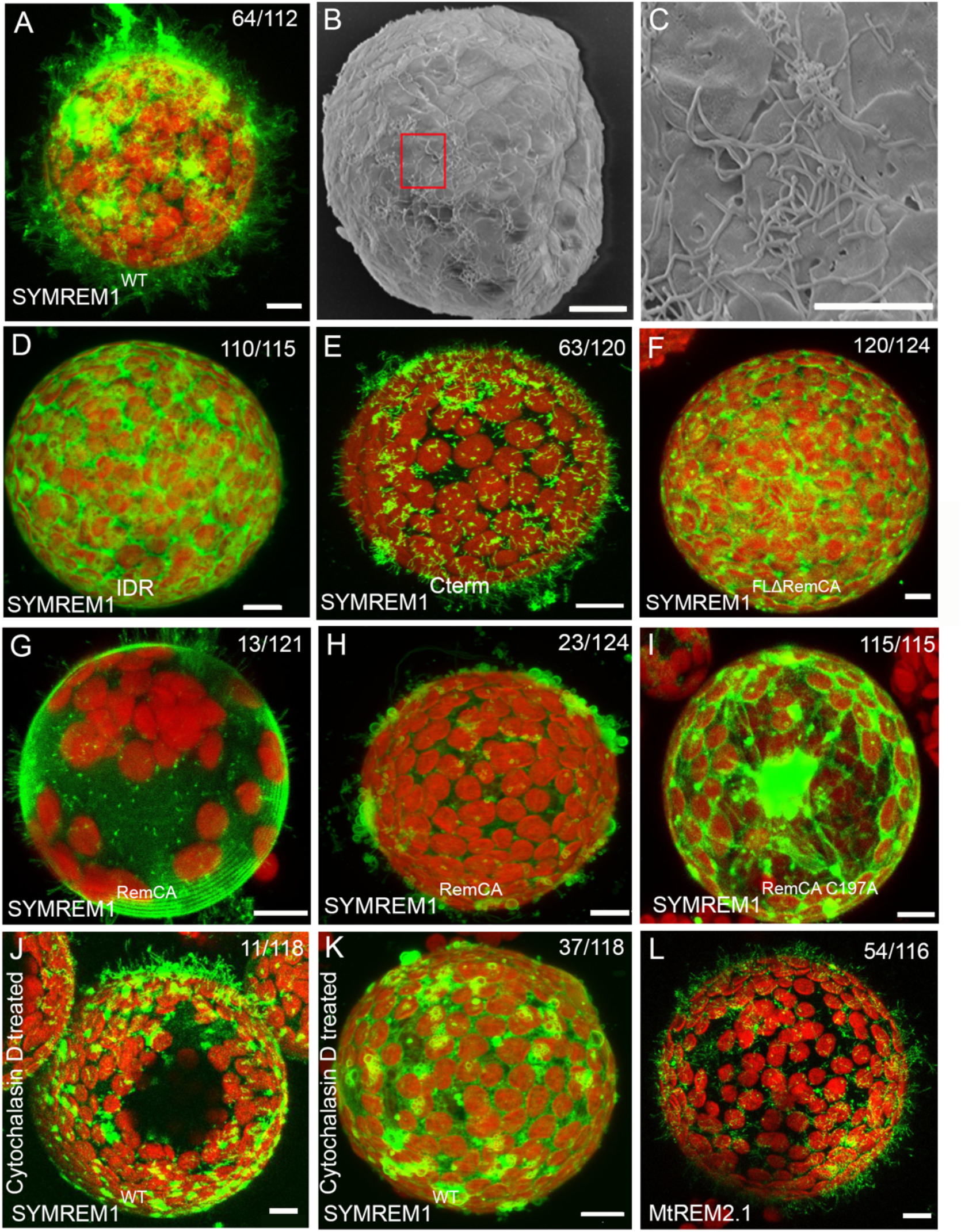
SYMREM1 stabilizes membrane tubulation and curvature in a cell wall-independent manner. *N. benthamiana* protoplasts ectopically expressing YFP-SYMREM1 develop multiple membrane tubes as shown by confocal laser-scanning (A) and scanning electron (B and close-up in C) microscopy within 2 hours after cell wall removal. (D-H) protoplasts expressing different SYMREM1 variants: only IDR domain (SYMREM1^IDR^, D), only C-terminal helix including RemCA (SYMREM1^Cterm^, E), full length sequence lacking the RemCA domain (F), RemCA domain only (G and H), and mutating the palmitoylated Cys197 residue within the RemCA domain (I). (J-K) Protoplasts ectopically expressing YFP-SYMREM1 after Cytochalasin D treatment. (L) *N. benthamiana* protoplasts ectopically expressing GFP-MtREM2.1 developed multiple membrane tubes. Scale bars indicate 10 μm. All the confocal images are shown as maximal projections. Red signals derive from chlorophyll autofluorescence within chloroplasts. Numbers indicate frequencies of observations.

Since SYMREM1 associates with the plasma membrane via the amphipathic and often palmitoylated RemCA peptide (amino acids 171-205) ^33^, we tested the impact of these C-terminal 35 amino acids. Truncating RemCA from full-length SYMREM1 (SYMREM1^FLΔRemCA^) resulted in a cytosolic protein that was unable to induce and/or stabilize these membrane outgrowths (Fig. 4F). Expression of the RemCA membrane anchor alone (SYMREM1^RemCA^) was sufficient to drive membrane topology changes. While 13 out of 121 protoplasts developed tubes on the protoplast surface (Fig. 4G), 23 out of 121 protoplasts exhibited less confined and more bulky membrane blebs (Fig. 4H). Mutating the palmitoylated Cys197 residue in RemCA (SYMREM1^RemCA C197A^) fully abolished membrane association of the peptide and its impact on membrane topology (Fig. 4I). These data show that the RemCA peptide alone is sufficient for initiating membrane tubulation but cannot maintain and/or stabilize long protrusions at high-frequency.

As tubular membrane outgrowths such as pollen tubes and root hairs are usually actin-dependent, we co-expressed full-length SYMREM1^WT^ with the actin marker LifeAct in protoplasts and observed that all tubes contained central actin filaments (Fig. 5A-A’’). The importance of this was additionally supported by the fact that application of cytochalasin D, an actin depolymerizing agent, dramatically decreased the number of tubular outgrowths (11 out of 118 protoplasts) in protoplasts expressing SYMREM1^WT^ (Fig. 4J). Instead, upon actin depolymerization we observed the same kind of membrane blebs as induced upon expression of the SYMREM1^RemCA^ peptide (Fig. 4K). Furthermore, expression of the symbiotic and actin-associated formin protein SYFO1 ^14^ resulted in a tip-localized signal at these protrusions (Fig. 5B-B’’) further implying that growth of these membrane tubes is, most likely, driven by actin prolongation. Since cytochalasin D treatment of SYMREM1^WT^ expressing protoplasts severely inhibited membrane tubulation we concluded that SYMREM1 stabilizes rather than actively drives outgrowth of these membrane tubes in an actin-dependent manner.

**Fig. 5:**
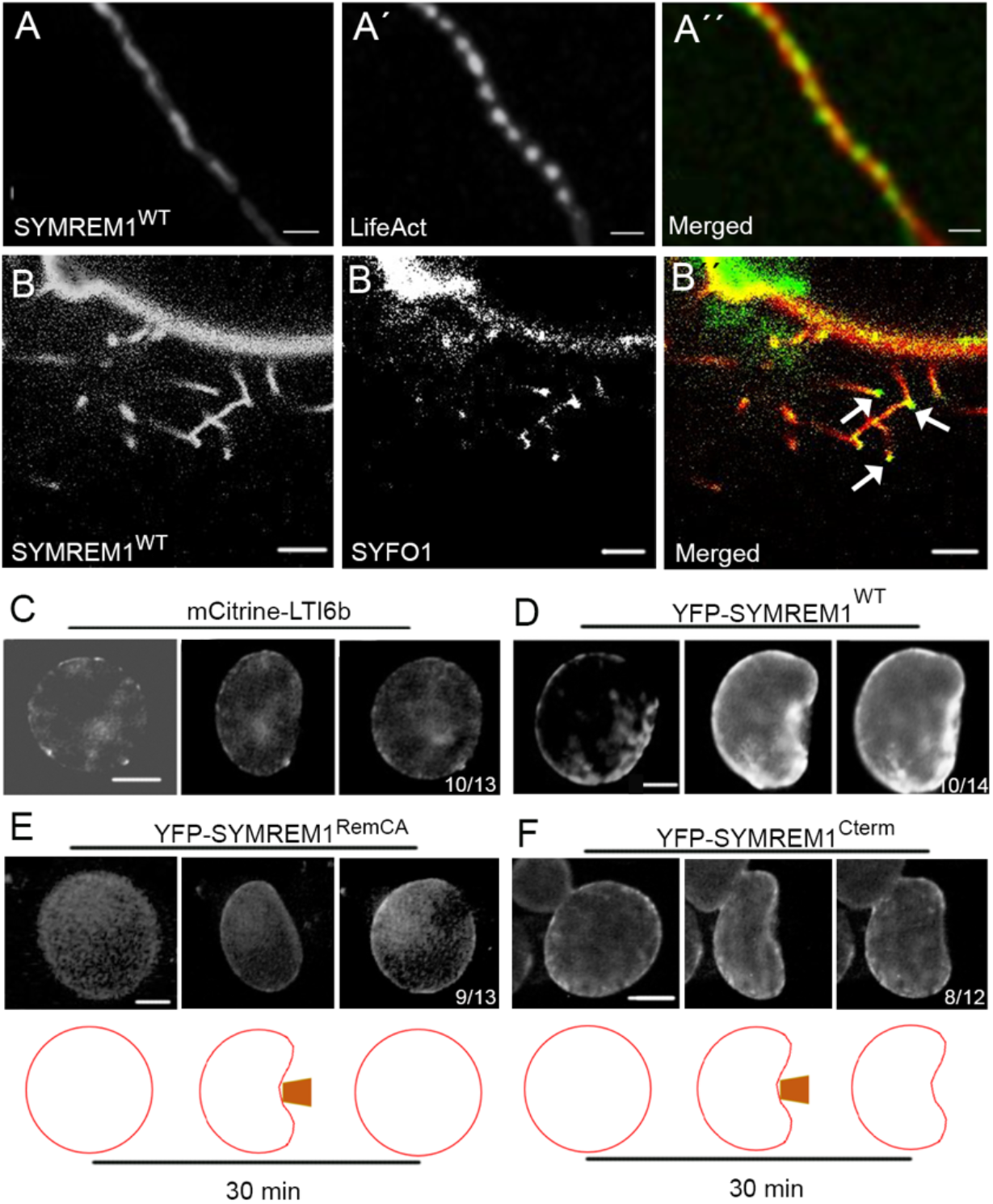
SYMREM1 stabilizes membrane negative membrane curvature in a cell wall-independent manner. *N. benthamiana* protoplasts ectopically expressing YFP-SYMREM1 developed multiple membrane tubes that comprised a central actin strand as labelled by LifeAct (A-A’’) with a tip localized formin protein SYFO1 (B-B’’, white arrows). Wall-less control protoplasts expressing the membrane marker mCitrine-LTI6b (C) and YFP-SYMREM1^RemCA^ (E) re-inflated immediately after 30 minutes of micro-capillary-based indentation, while the great majority of those expressing YFP-SYMREM1^WT^ (D) and YFP-SYMREM1^Cterm^ (F) retained the induced membrane curvature. Scale bars indicate 15 μm (A) 10 μm (B, C-D’’), 5 μm (B’) and 25 μm (E-F).

To test whether SYMREM1 functions as a stabilizing scaffold exclusively for positive curvatures, we assessed whether negative curvatures, as underlying IT-associated membrane tubes, were also maintained in the presence of this protein. Therefore, we isolated SYMREM1-expressing protoplasts and immediately indented them with a micro-capillary for 30 minutes. Here, 10/13 protoplasts expressing the LTI6b membrane marker (control) re-inflated directly after releasing the pressure (Fig. 5C) indicating that microtubules alone are not sufficient to dynamically maintain these short-term indentations as shown for protoplasts that had been confined in controlled geometries^45^. By contrast, full re-inflation was observed in only 4/14 protoplasts expressing SYMREM1, while microcapillary-induced membrane deformations were maintained in the majority (10/14) of these protoplasts (Fig. 5D). Such stabilization was also observed in most protoplasts expressing the truncated SYMREM1^Cterm^ variant (Fig. 5F) while the RemCA peptide alone (SYMREM1^RemCA^) was not sufficient to mediate this phenomenon (Fig. 5E). These data further support the role of SYMREM1 as a membrane topology scaffold.

As a last set of experiments, we assessed whether membrane topology scaffolding is limited to SYMREM1 or a more wide-spread feature within the remorin protein family. In analogy to the Medicago SYMREM1 (MtSYMREM1) protein, expression of the orthologous gene from *Lotus japonicus* (*LjSYMREM1*; Lj4g3v2928720) resulted in stabilized membrane tubes at high frequency (Fig. S4A). Evolutionary, the presence of SYMREM1 proteins is mainly restricted to species that maintained the root nodule symbiosis and species that are not able to allow intracellular infection of symbionts, either fungal or bacterial, through their epidermis have lost *SYMREM 1* (Fig. S5). In addition, the only non-symbiotic clade that retained *SYMREM1*, the Caryophyllales, experienced bursts of diversifying positive selection indicative of neofunctionalization (Table S1), suggesting that purifying selection maintaining *SYMREM1* in plant genomes is linked to the maintenance of symbiotic abilities. Interestingly, a closely-related protein (MedtruChr5g0397781; MtREM2.1) shows an expanded phylogenic distribution, while both *SYMREM1* and *MtREM2*.*1* derive from a Papilionoideae duplication. To assess whether MtREM2.1 has similar impact on membrane topology as shown for SYMREM1, we expressed a MtREM2.1 fusion protein in mesophyll protoplasts. Indeed, the presence of this protein resulted in the stabilization of numerous membrane tubes (Fig. 4L), and this function is supported by AlphaFold predictions that revealed a putative structure that is highly similar to the one predicted for SYMREM1 (Fig. 3D-F). To address whether MtREM2.1 localizes to highly curved membranes, we first surveyed its gene expression profile by using global transcriptome data, which revealed an induced expression during endosymbiotic interactions ^46^. We independently confirmed this by quantitative RealTime PCR on cDNA generated from nodulated and mycorrhized roots (Fig. S6A). To increase the spatio-temporal resolution of this analysis, we generated a transcriptional reporter with a 1kb long fragment of the putative *MtREM2*.*1* promoter driving the expression of a β-glucuronidase gene (GUS). *MtREM2*.*1* promoter activity was detected in young and maintained in the apical region of mature nodules (Fig. S6B-6C’). In addition, we found increased GUS activity in cells containing arbuscules when inoculating roots with the arbuscular mycorrhizal fungus *Rhizophagus irregularis* (Fig. S6E-6F’). To assess the localization of the MtREM2.1 protein, we generated *Medicago truncatula* stable lines (WT R108) expressing an mCherry-MtREM2.1 fusion proteins under the control of the endogenous promoter. In roots inoculated with *R. irregularis*, MtREM2.1 predominantly localized to the host-derived and highly curved membrane (periarbuscular membrane) surrounding intracellular hyphae, from their early penetration into inner cortical cells (Fig. S6G-6G’) until development of fully branched arbuscules (Fig. S6JH-S6H’). Interestingly this arbuscular membrane is, similarly to infection droplets or symbiosome membranes, not supported by a rigid cell wall ^47^.

While all three tested group 2 remorins (MtSYMREM1, LjSYMREM1 and MtREM2.1) stabilized membrane tubes on protoplasts, systematic expression of different members of the Arabidopsis remorin family, confirmed this ability for AtREM1.3 (Fig. S4B), AtREM1.4 (Fig. S4D), and AtREM4.2 (Fig. S4G) even though membrane tubulation was stabilized with much lower frequencies in all cases. In addition, and similar to the effect observed in the presence of RemCA (Fig. 4H) or cytochalasin D (Fig. 4K), expression of AtREM1.3 (Fig. S4C) resulted in membrane bleb formation in 11% of the cases. No such effects were observed for AtREM2.1 (Fig. S4E), which was mostly cytosolic, whereas expression of AtREM3.2 (Fig. S4F), AtREM5.1 (Fig. S4H) and AtREM6.1 (Fig. S4I) led to focal accumulations and the formation of shorter membrane tubes. These results indicate that the ability of remorins to act as membrane topology scaffolds is conserved in land plants. Besides this, it was intriguing to note that expression of an N-terminally truncated variant of AtREM1.3 lacking its IDR significantly increased the ability to stabilize membrane tubes from 6% (Fig. S4B) to 24% (Fig. S4J), suggesting an inhibitory role of the disordered N-terminal region. This hypothesis was supported by expressing a chimeric protein comprised of the AtREM1.3 N-terminal and the MtSYMREM1 C-terminal region (AtREM1.3^N-term^::MtSYMREM1^C-term^) that reduced membrane tubulation frequencies from 57% (Fig. 4A) to 17% (Fig. S4K), while expression of a LjSYMREM1/MtSYMREM1 chimera maintained high tubulation levels at 47% (Fig. S4L). Taken together, these results show that the largely legume-confined remorins SYMREM1 and MtREM2.1 represent evolutionary optimizations to structurally support curvatures of cell wall-devoid membranes throughout endosymbiotic infection processes.

## Discussion

Plant membranes are usually supported by a rigid cell wall that counteracts the intracellular turgor pressure and maintains, together with cortical microtubules and actin, cellular shapes over their lifespan ^45,48-50^. As all cell wall constituents are either secreted to the apoplast or synthesized directly onto the extracellular leaflet of the plasma membrane, these responses require time. Furthermore, subsequent loosening of the rigidified cell wall to adopt temporal changes in membrane topologies mostly account for protrusion-like shapes. In contrast, larger scale and negatively curved membrane invaginations as found during microbial infections of host cells may require other types of scaffolding prior to de novo cell wall apposition. Such functions can be maintained by scaffold proteins as exemplified during clathrin-mediated endocytosis, where the adaptor complex AP2 together with clathrin light and heavy chains stabilizes endocytotic vesicles ^22,23^. In plants, evidence has been presented that endocytosis of membrane nanodomain-localized proteins might occur via a flotillin- and remorin-dependent but clathrin-independent pathway ^51-53^. However, compelling evidence for topological scaffolding functions of these proteins has been missing, while the impact of remorins on membrane fluidity is experimentally supported ^28^.

Here, we demonstrate that plant remorin proteins can stabilize membrane topologies independent of a cell wall with group 2 remorins, which are mostly found in plants that have maintained a symbiotic association with (both) rhizobia and arbuscular mycorrhiza fungi, showing the strongest effect on membrane tubulation (Fig. 4, Fig. S4). This makes sense since both endosymbioses maintain highly curved perimicrobial membranes such as the periarbuscular and the IT membrane. Although cell wall components are deposited within the older parts of ITs ^20^, the initial curvature needs to be differently stabilized and additionally maintained around cell wall-free bacterial release sites. Using microcapillary-generated forces, we demonstrate that the evolutionary conserved C-terminal region of the remorin SYMREM1 is able to stabilize such negatively curved membrane invaginations (Fig. 5F). This segment of the protein itself is required for remorin oligomerization into dimers via hydrogen bonding and hydrophobic interactions (Fig. 3). These higher order oligomers can also be found in SYMREM1-stabilized membrane tubes *in vivo* where it may wrap around actin filaments in a helical arrangement (Fig. 5A). Similar observations have been made for the group 1 remorin REM1.3 from *Arabidopsis thaliana* when being expressed in human Cos-7 cells ^54^. Tight links of remorin to actin are also supported by SYMREM1 colocalization with the formin protein SYFO1 in protoplast tubes (Fig. 5B), the control of formin condensation by remorins during innate immune responses in Arabidopsis ^35^, the interaction of the group 6 remorin GSD1 from rice with actin ^55^ and the fact that actin densely accumulates, as also shown for SYMREM1 (Fig. 2), at infection droplets ^56^. Additional evidence for remorins being able to form higher order structures also comes from *in vitro* experiments, where the C-terminal region of a group 1 remorin auto-assembles into proteinaceous sheets ^31^, further supporting the hypothesis that remorins function as topological membrane scaffolds. The impact of such protein lattices on membrane curvature has also been demonstrated for the human N-Bar protein Endophilin that can adopt higher-order arrangements ^57^. Similar observations have also reported for the F-Bar protein Imp2 from fission yeast, where a helical alignment of Imp2-subunits results in membrane tubulation of human Cos-7 cells ^58^, a feature that was also reported upon expression of the Arabidopsis remorin REM1.3 in this cell line ^54^.

While these effects are mediated by the antiparallel dimers of the C-terminal α-helices or their higher order oligomers, the N-terminal IDR most likely serves regulatory functions as a chimeric protein that contains the structured C-terminal region of the Medicago SYMREM1 protein but the IDR of the Arabidopsis REM1.3 exhibits a reduced ability to stabilize membrane tubulation in protoplasts (Fig. S4K). However, this feature is not compromised when using the N-terminal IDR from the orthologous Lotus SYMREM1 protein (Fig. S4L). A conservation to induce efficient membrane tubulation among symbiotically active remorins is also in line with the observation that *symrem1* mutants show defects in bacterial infection and release, which might be linked to the inability to temporarily stabilize secondary membrane invaginations as precursors of bacterial release sites (Fig. 2D-2M). Instead, cells of *symrem1* mutants that have been passed by an IT contain a multitude of empty and unsupported membrane spheres (Fig. 1C-1H).

The fact that remorins associate with the inner leaflet of the plasma membrane by their amphipathic C-terminal anchor peptide (RemCA) and palmitoylation ^27,33,34^ and that this process impacts membrane fluidity ^28^, the condensation of formin ^35^ and the microtubule binding protein DREPP ^59^ support the hypothesis that remorins also drive targeted secretion of proteins and lipids. This together with the intrinsic ability to form oligomeric structures might explain that these proteins can stabilize positive and negative curvatures as observed in plasmodesmata ^29,54^, the extrahaustorial membrane ^60^, the symbiosome membrane ^36,43^ and the periarbuscular membrane (Fig. S6G-H’). It would indeed be challenging to test, whether the stabilization of type II intra-matrix compartments, which are swirl-like extensions from the periarbuscular membrane into the periarbuscular space ^61,62^, is remorin-dependent.

Taken together we unraveled a novel mechanism that allows plant cells to stabilize distinct membrane topologies in the absence of a cell wall by an interplay between oligomeric remorin scaffolds and the cytoskeleton. Consequently, remorins may serve additional functions that have been mainly attributed to BAR-domain proteins in mammalian cells ^63^.

## Materials and Methods

### Plant growth and Rhizobia inoculation

Seeds of *M. truncatula* were surface sterilized by covering them with pure sulfuric acid (H_2_SO_4_) for 10 minutes, followed by 4 to 6 times washing with sterile water. Seeds were then covered with bleaching solution (12% NaOCl, 0.1% SDS) for no longer than 60 seconds and washed 4 to 6 times with sterile water. After surface sterilization, seeds were transferred to 1% agar plates and stratified at 4°C for 3 days in darkness. Germination was allowed for up to 24 hours at 24°C in darkness. The seed coat was removed and the seedlings were transferred to plates containing Fahräeus medium supplemented with 0.5 mM NH_4_NO_3_. One week later, they were transferred onto new Fahräeus medium without nitrate, but containing AVG (0.1 μM). Inoculations were performed after 4 days of growth on plates without nitrate. For inoculation of *M. truncatula* roots, a *S. meliloti* (Sm2011) liquid culture was centrifuged (3 min, 3000 rpm), washed once with liquid Fahräeus medium and resuspended in liquid Fahräeus medium to a final OD_600_ = 0.03. Each root was covered with 1 ml of rhizobia suspension, which was removed after 6 minutes. Afterwards, the plants were placed in a controlled environment chamber at 24°C with a 16/8 hours light/dark photoperiod, keeping the roots in the dark, for 3 weeks before harvesting the nodules.

### Hairy root transformation and Rhizobia inoculation

*M. truncatula* hairy root transformation was performed as previously described ^64^. Briefly, transgenic *Agrobacterium rhizogenes* (ARqua I), carrying the plasmid of interest, was grown in LB liquid culture for one day and 300 μl of the culture were spread on LB agar plates supplemented with the corresponding antibiotics for selection, and incubated on plates for two more days before transformation. *M. truncatula* seeds were prepared as mentioned above. After germination the seed coat was removed from the cotyledons of the seedlings under water, and the root meristem was cut off with a scalpel. Cut seedlings were dipped on the Agrobacterium plates and transferred onto solid Fahräeus medium (containing 0.5 mM NH_4_NO_3_). Transformed seedlings were incubated for three days at 22°C in darkness, following 4 days at 22°C in white light but keeping the roots in the dark. One week after transformation, seedlings were transferred onto new Fahräeus medium (0.5 mM NH_4_NO_3_) and grown for another 10 days at 24°C in a controlled environment chamber with 16h/8 hours light/dark photoperiod. Afterwards, the roots were screened to examine the transformation efficiency using a stereomicroscope to detect the corresponding fluorescent signal. Untransformed roots were cut off and plants showing fluorescence roots were transferred to pots (2 plants per pot) having a mixture of equal volume of quarzsand and vermiculite. All pots were individually watered with liquid Fahräeus medium (without nitrate) and tap water once a week. After 3-5 days, the pots were inoculated with *S. meliloti* (OD_600_ = 0.003).

### Evolutionary analysis

Homologs of MtSYMREM1 (Medtr8g097320.2 or MtrunA17Chr8g0386521) were retrieved using the tBLASTn v2.11.0+ ^65^ against a database of 189 species covering all Viridiplantae lineages with default parameters and e-value threshold of 1e-10. Coding sequences of putative homologs were aligned using MAFFT v7.471 ^66^ with default parameters. Positions with more than 80% of gaps were removed from the subsequent alignment using trimAl v1.4rev15 ^67^ and cleaned alignment subjected to phylogenetic analysis using Maximum Likelihood approach. Prior tree reconstruction, best-fitting evolutionary model was tested using ModelFinder ^68^ according to the Bayesian Information Criteria. Maximum likelihood analysis was conducted using IQ-TREE2 v2.0.3 ^69^. Branches support were estimated using 10,000 replicates of both sh-aLRT ^70^ and UltraFast Bootstraps ^71^. Analysis of the main tree revealed that MtSYMREM1 clade derives from Eudicots duplication and then, a subtree corresponding to the Eudicots clade of MtSYMREM1 has been reconstructed based on protein sequences aligned using MUSCLE v3.8.1551 ^72^ and phylogenetic procedure described above. Trees were visualized using the iTOL v6 platform ^73^.

To look for specific selective pressure acting on the Caryophyllales clade, which is composed only of species not forming infection threads, we conducted branch and branch site analysis. We estimated relaxation (K<1) or intensification (K>1) parameters and also positive selection acting on Caryphyllales (Table S1), we used the RELAX and aBSREL programs implemented in the HYHPY software ^74,75^. These methods calculate different synonymous and nonsynonymous substitution rates (ω = *d*N/*d*S) using the phylogenetic tree topology for both foreground and background branches. Translated CDS of MtSYMREM1 homologs were aligned using DECIPHER ^76^. In total, 191 sequences and 162 codons were analysed (Table S2).

### Nodule sections

For analysing the subcellular localization of SYMREM1 and the LactC2 biosensor in WT and *ipd3* (*sym1-TE7*) mutant plants, the corresponding constructs were used to transform Medicago plants by hairy root transformation as described above. Nodules were harvested 2 weeks after being inoculated with *S. meliloti* (mCherry) in open pots and directly embedded in 7 % low melting agarose. Semi-thin (70μm) longitudinal sections were obtained using a vibratome microtome (VT1000S, Leica) and the sections were analyzed using a confocal microscope (Leica TCS SP8).

### Construct Design

Constructs used for SYMREM1 localization assays, protoplast analyses, and yeast transformation are described in ^33^. For *Symrem1* expression and purification, the *Symrem1* coding sequence of *Medicago truncatula* was recombined into the Gateway (GW) compatible pDEST17 vector via LR-reaction. The PS reporter (2x LactC2 domain), the plasma membrane marker (LTI6b), and all the chimeric sequences were synthesized by Life Technologies, then cloned to expression vector by Golden Gate cloning ^77^. All the designed constructs and all primers used are listed in Table S3 and Table S4. The sequence data from this article can be found in phytozome (https://phytozome.jgi.doe.gov/) with the gene ID: *Symrem1*(Medtr8g097320), *Syfo1*(Medtr5g036540).

For promoter activation studies, a sequence 1 kb upstream of the start codon of the *MtRem2*.*1* gene was amplified via PCR from genomic DNA (A17) and cloned into a pENTR/D-TOPO vector. The *ProMycrem*:eGFP-GUS construct was created by a LR-reaction of pKGWFS7-vector and pENTR/D-TOPO: *ProMycrem*.

### Histochemical Promoter analysis (GUS-staining), WGA staining and Microscopy

The activation patterns of the *MtRem2*.*1* promoter were analyzed via β -Glucuronidase (GUS) activity. Transgenic roots were stained in GUS-staining solution (0.1 M NaPO_4_; 1 mM EDTA; 1 mM K_3_Fe (CN)_6_; 1 mM K_4_Fe (CN)_6_; 1% Triton-X 100;1 mM X-Gluc) at 37°C for 4 hours in the dark. For fluorescent visualization of fungal structures, colonized roots were fixed in 50% ethanol for at least 12 hours and afterwards cleared for 2 days at room temperature in 10% KOH. After a washing step with distilled water, roots were incubated in 0.1 M HCl for 1 hour at RT. Prior the final staining, roots were washed with distilled water and rinsed once with PBS (phosphate buffered saline; pH7.4). Roots were placed in a PBS-WGA–AlexaFluor594 staining solution (0.2 μg/mL WGA-AlexaFluor594; Thermo Fisher Scientific, Germany) for at least 6 hours at 4°C in dark.

### Expression analysis

Total RNA extraction was performed according to the Spectrum Plant Total RNA Kit (Sigma-Aldrich, Germany) manual. Root material was grinded in liquid nitrogen and 100 mg per root sample was used for extraction. Extracted RNA was treated with DNase I, Amp Grade (Invitrogen, Germany). The absence of genomic DNA was verified via PCR. Synthesis of cDNA was performed with 700 ng of RNA in a total reaction volume of 20 μl using the Superscript III kit (Invitrogen, Germany). For qRT-PCR analysis, a Fast SYBR Green Master Mix (Applied Biosystems, Gemany) was used in a 10 μl reaction volume. A CFX96TM Real-Time system (Bio-Rad, Germany) was used for PCR reactions and detection. Expression was normalized to *Ubiquitin*. At least three biological replicates were analysed in technical duplicates per treatment.

### Transformation of *Nicotiana benthamiana* leaves and protoplast isolation

Transgenic *Agrobacterium tumefaciens* carrying the plasmids of interest were grown in LB liquid culture overnight at 28°C with the appropriate antibiotics. The culture was centrifuged (4,000 rpm, 2 min) and the pellet was resuspended in Agromix (10 mM MgCl_2_; 10 mM MES/KOH pH 5.6; 150 μM Acetosyringone) to an OD600 of 0.3. Bacteria were mixed with the silencing suppressor p19 before being incubated for 2 hours at 25°C in darkness and then infiltrated at the lower site of *N. benthamiana* leaves. Two days after infiltration, the transformed leaves were harvested to isolate protoplasts according as described earlier ^78^ with a small modification (PNT solution contained 300 mg/l of CaCl_2_ 2H_2_O). All experiments using protoplasts were done at least three times independently.

### Microcapillary assay

Isolated protoplasts were embedded in 0.5% agarose on a cover of a Petri Dish. The injection set-up consisted of an inverted microscope (Zeiss Axiovert 135 TV) with a motor driven micromanipulator (LANG GmbH & Co. KG, Type: STM3) mounted at the right side of the stage. Femtotips injection needles (Eppendorf) were adapted by removing the sharp-pointed tip of the needle by hand, until obtaining a needle that could not penetrate the protoplast plasma-membrane.

### Confocal Laser-Scanning Microscopy

Protoplast images were obtained using a Leica TCS SP8 confocal microscope equipped with a 20x water immersion lens (Leica Microsystems, Mannheim, Germany), with the exception that images from Fig 2A and Fig 4A-A’’ were taken with a Zeiss LSM880 Airyscan using a 63x oil immersion lens. GFP was excited with a White Light Laser (WLL) at 488 nm and the emission detected at 500-550 nm. YFP was excited with a 514 nm laser line and detected at 520–555nm. mCherry fluorescence was excited at 561 nm and emission was detected between 575-630 nm. Samples, co-expressing two fluorophores were imaged in sequential mode between frames. All image analyses and projections were performed with either ImageJ/(Fiji) software ^79^ or Imaris.

### Protein expression and purification

*E. coli* BL21(DE3) cells were transformed with plasmid pDEST17 encoding His-SYMREM1 protein. A single colony of transformed *E. coli* was transferred in LB medium and grown overnight for obtaining a pre-culture. Then, 40 ml of pre-culture was inoculated in 2 l of LB media and further grown at 37°C. Protein expression was induced by 1 mM IPTG at OD_600_ of 0.6. Afterwards, cells were incubated overnight (about 20 hours) at 25°C. Cells were harvested by centrifugation at 6000g for 15 min. The cell pellet was resuspended in 100 ml Lysis buffer (20 mM HEPES, 500 mM NaCl, 20 mM imidazole, 10 % glycerol, 1 mM EDTA, 1 mM Pefabloc, pH 7.2) and cells were passed through Constant Cell Disrupter (Constant Systems Limited). Cell debris was removed by centrifugation at 30,000g for 30 min. The cleared cell lysate was loaded onto IMAC column (5 ml HisTrap_FF) pre-equilibrated with Loading buffer A (20 mM HEPES, 500 mM NaCl, 20 mM imidazole, pH 7.2) and washed with 10 CV of Loading buffer A. Proteins were eluted with a linear gradient of imidazole from 20 to 450 mM in 15 CV. The eluted fractions were pooled and concentrated by spin filtration to 5 ml. Precipitated proteins were removed by an additional centrifugation for 10 min at 10,000g before loading onto gel-filtration column (HiLoad Superdex 200 16/60) equilibrated with PBS. Eluted fractions after gel-filtration were analysed with SDS-PAGE. Those fractions containing pure His-SYMREM1 were pooled and concentrated by spin filtration to the working concentration. For Immunoblot Analysis: 2 weeks old nodules were harvested. Extracted proteins were separated on a 12% (w/v) SDS-PAGE gel and transferred overnight at 30 V to a PVDF (polyvinylidene difluoride) membrane. Membranes were then blocked with 5% (w/v) milk for 1 hour at room temperature before being hybridized with the SYMREM1 peptide antibodies ^36^ at a dilution of 1:500 for 1.5 hours at room temperature. Membranes were then washed with TBST three times for 10min before incubating with a second antibody (anti-rabbit (Sigma), 1:2000 dilution, 1 hour at room temperature). Prior to image acquisition, the membranes were washed again with TBST three time for 10 min.

### Electron Microscopy

Transmission Electron Microscopy (TEM) was performed on nodules harvested at (3 wpi). Nodules were cut longitudinally in half, immediately fixed in MTSB buffer ^80^ containing 2.5% glutaraldehyde and 4% p-Formaldehyde under vacuum for 15 min (twice), and stored at 4°C in fixative solution until used further. After washing 5 times for 10 min each with buffer, nodules were post-fixed with 1% OsO_4_ in H_2_O at 4°C for 2 hours and again washed 5 times (10 min each) with H_2_O at room temperature. The tissue was in block stained with 1% Uranyl Acetate for 1 hour in darkness, washed 3 times (10 min each) in H_2_O, and dehydrated in EtOH/H_2_O graded series (30%, 50%, 70%, 80%, 90%, 95% 15 min each). Final dehydration was achieved by incubating the samples twice in absolute EtOH (30 min each) followed by incubation in dehydrated acetone twice (30 min each). Embedding of the samples was performed by gradually infiltrating them with Epoxy resin (Agar 100) mixed with acetone at 1:3, 1:1 and 3:1 ratio for 12 hours each, and finally in pure Epoxy resin for 48 hours with resin changes every 12 hours. Polymerization was carried out at 60°C for 48 hours. Ultrathin sections of approximately 70 nm were obtained with a Reichert-Jung ultra-microtome and collected in TEM slot grids. Images were acquired with a Philips CM 10 transmission electron microscope coupled to a Gatan BioScan 792 CCD camera at 80 kV acceleration voltage.

Scanning Electron Microscopy was carried out on freshly isolated protoplasts and on longitudinal vibratome sections (70μm) of nodules collected after 3 wpi. The material was immediately fixed, dehydrated in graded EtOH series until 100% EtOH, and critical point dried in absolute EtOH-CO_2_. Dried material was mounted on carbon tabs and coated with 5 nm platinum. Imaging of samples was performed using a Hitachi S-4800 microscope at 5kV acceleration voltage.

Negative staining of purified SYMREM1 protein was performed by applying 5 *μ*l protein solution to glow-discharged 400 Cu mesh carbon grids for 10 min, blotting and negatively staining using 2% (w/v) uranyl acetate. Images were recorded under low-dose conditions on a Talos F200C transmission electron microscope operated at 200 kV and equipped with a Ceta 16M camera. Micrographs were taken at a nominal magnification of 73,000x. A total of 389 segments were manually selected using RELION-3.1.0 ^81^. The defocus and astigmatism of the images were determined with CTFFIND4.1 ^82^ and numerical phase-flipping was done to correct for effects of the contrast transfer function using RELION-3.1.0. Image processing was done using IMAGIC-5 ^83^. Particle images were band pass filtered between 400 and 10 Å, normalized and centred by iteratively aligning them to a vertically oriented class average. Class averages containing 5-10 images were obtained by four rounds of classification based on multivariate statistical analysis, followed by multi-reference alignment using homogenous classes as new references.

### Protein 3D predication

In-silico structure predictions were made with DeepMind AlphaFold 2.1.1 in multimer mode ^41,42^. Electrostatic surface potentials were generated with ABPS ^84^, figures were created with PyMOL (Schrödinger LLC).

## Acknowledgements

We would like to thank Carmen Schubert and Rosula Hinnenberg for their excellent technical help and the entire Ott lab team for fruitful discussions and providing their individual expertise throughout the course of the project. We also thank the staff of the Life Imaging Center (LIC) in the Hilde Mangold House (HMH) of the Albert-Ludwigs-University of Freiburg for help with their confocal microscopy resources, and the excellent support in image recording. Special thanks also to Norbert Roos and Jens Wohlmann at the Electron Microscopy Facility, Department of Biosciences, University of Oslo, Norway, for helping with the SEM sample preparation and imaging and to Falk Tauber (University of Freiburg, Cluster of Excellence livMathS) for designing and printing a tentative indentation support. Many thanks also to Julien Gronnier (University of Tübingen) for providing the REM2.1 CDS and for critically reading the manuscript. JK, CL and PMD belong to the LRSV, which is part of the TULIP LABEX (ANR-10-LABX-41). The microscopes are operated by the Microscopy and Image Analysis Platform (MIAP) and the Life Imaging Center (LIC), Freiburg.

## Funding

Engineering Nitrogen Symbiosis for Africa (ENSA) project currently supported through a grant to the University of Cambridge by the Bill & Melinda Gates Foundation (OPP1172165) and UK government’s Department for International Development (DFID) (TO, PMD)

Deutsche Forschungsgemeinschaft (DFG, German Research Foundation) 431626755 (TO), 442219341 (PW)

DFG under Germany’s Excellence Strategy grant CIBSS – EXC-2189 – Project ID 39093984 (TO/CH/WW)

China Scholarship Council (CSC) grants 201708080016 and 201506350004 (CS/PL)

DFG project number 414136422 (CLSM; TO), DFG project number 426849454 (TEM; TO) and DFG project number 406260942 (cryo-TEM; PW)

## Author Contributions

Conceptualization, CS and TO; Investigation, CS, MR-F, BL, CHR, NN, AAMF, ES, KEG, EVM, NG, EM, CP, PL, JK, CL, PW, TS, PMD, OE and TO; Writing –Original Draft, CS, MR-F, and TO; Writing –Review & Editing, CS, MR-F, BL, CHR, PL, NN, AAMF, ES, KEG, EM, NG, EM, JK, CL, PMD, CH, WW, PW, TS, OE and TO; Funding Acquisition, PL, CS, RG, PMD, PW, OE and TO; Supervision, PMD, CH, WW, PW, TS and TO.

## Data and materials availability

All data are available in the main text or the supplementary materials.

## Supplementary Materials

Figs. S1 to S6

Tables S1-S4

