## Supplemental Figures for "Remorin proteins serves as membrane topology scaffolds in plants"

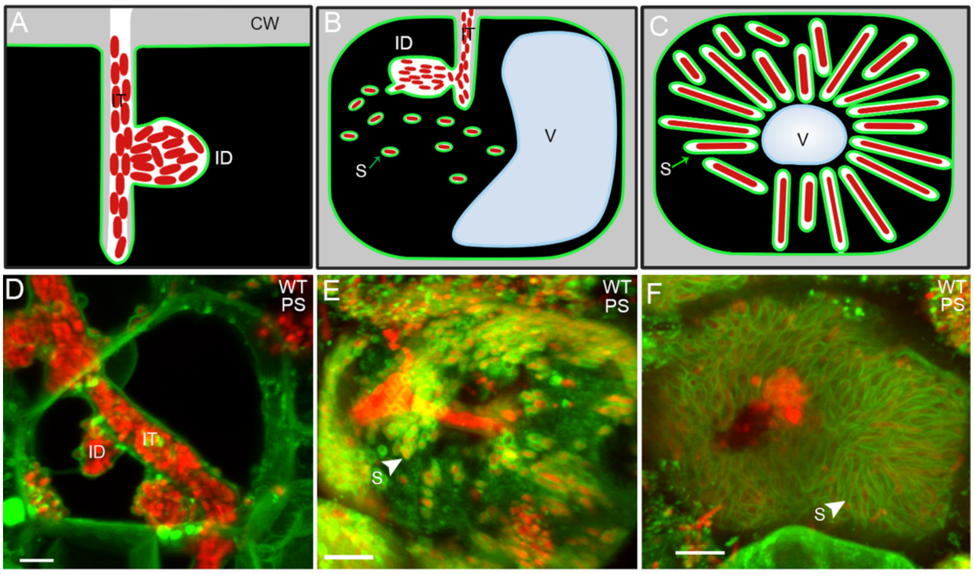


**Fig. S1. Visualization of symbiotic membranes using the phosphatidylserine reporter LactC2**. (A-C) Sketches of infected cells inside nodules at different stages (also used in Fig. 2). CW: cell wall; IT: infection thread; ID: infection droplet; S: symbiosome; V: vacuole. (D-F) To visualize membrane structures, phosphatidylserine (PS) was labelled using a LactC2 biosensor (*S. meliloti* expressing an mCherry marker; red). WT: Wild-type (A17). Arrow heads indicate symbiosomes (E, F). Scale bars indicate 5 µm (D-F).

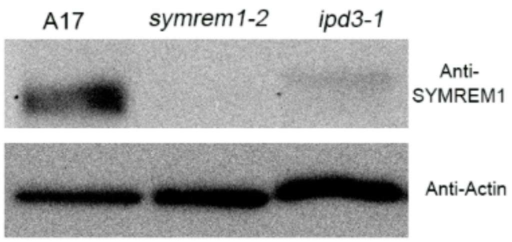


**Fig. S2. SYMREM1 protein levels are greatly reduced in *ipd3-1* mutant.** Nodules from *symrem1-2* and *ipd3-1* mutant plants were harvested at 2 weeks post-inoculation with *S. meliloti* and used for total protein extraction. Western blot analysis was performed using custom-made anti-SYMREM1 peptide and anti-actin antibodies.

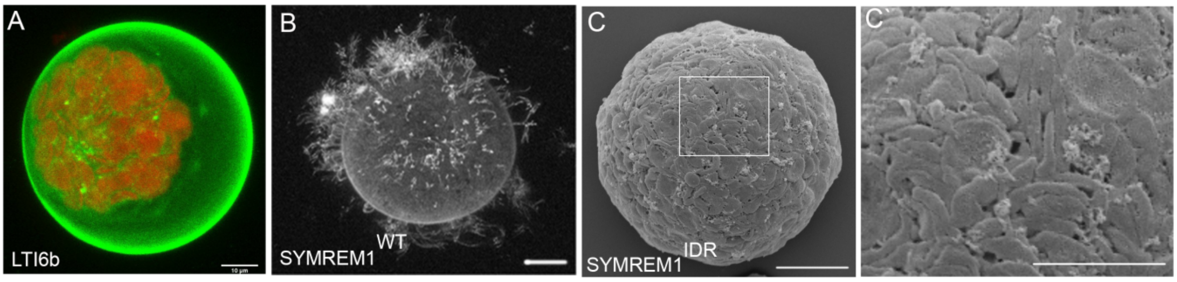


**Fig. S3. SYMREM1-induced membrane tubulation on protoplast surfaces.** Protoplasts expressing the control transmembrane domain of LTI6b maintained a smooth surface spherical shape (A). By contrast, strong membrane tubulation was observed on protoplasts expressing full-length SYMREM1 (B), while this was not observed upon expressing the isolated IDR segment (amino acids 1-73) of SYMREM1 (C, C`). (A-B) are maximum projections of confocal images, (C-D) images derived from scanning electron microscopy. Scale bars indicate 10 µm in (A, B, and C`) 20 µm in (C).

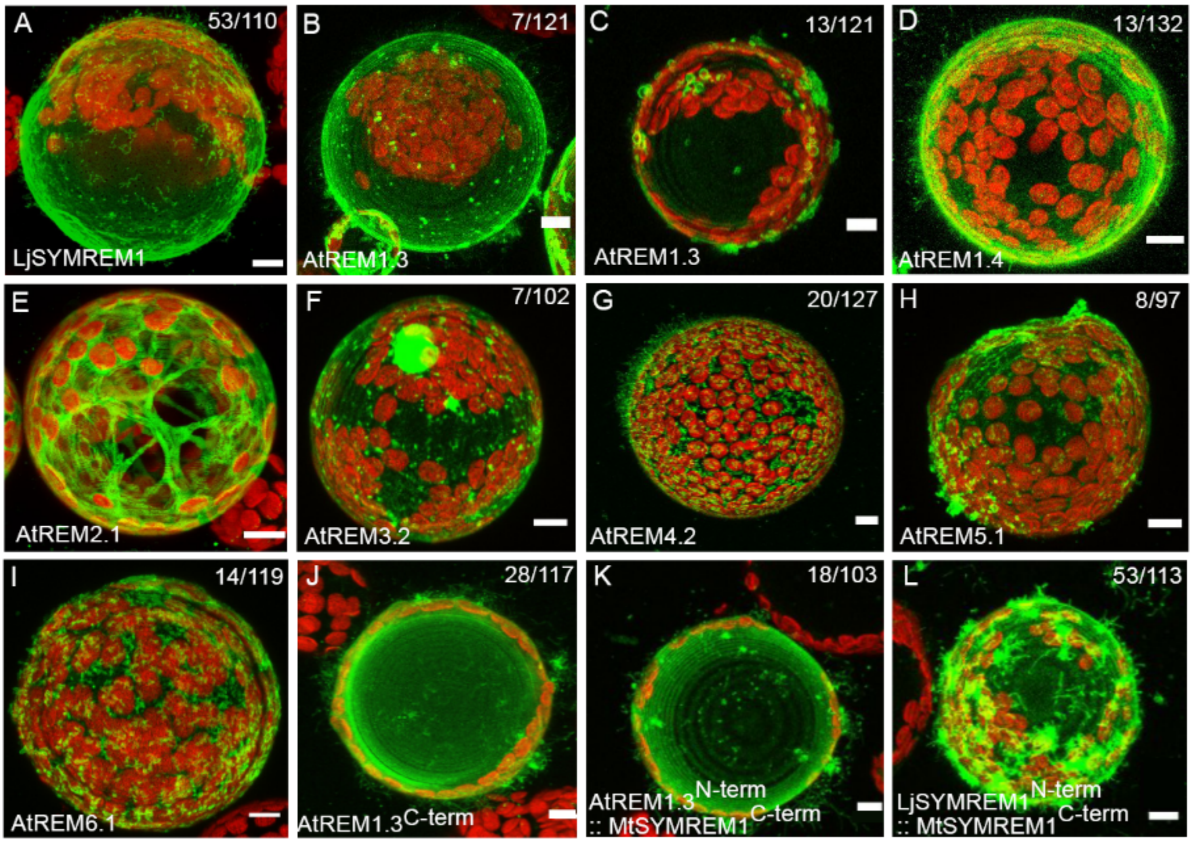


**Fig. S4. Remorin-induced membrane tubulation in protoplasts.** *N. benthamiana* protoplasts ectopically expressing Lotus SYMREM1 (A) and six different remorins from the non-symbiotic plant *Arabidopsis thaliana* covering all five sub-groups of the Arabidopsis remorin family (B-I). AtREM1.3 (B and C), AtREM1.4 (D), AtREM2.1 (E), AtREM3.2 (F), AtREM4.2 (G), AtREM5.1 (H), and AtREM6.1 (Iwere expressed as N-terminally tagged YFP fusion proteins for at least 2 hours prior to confocal imaging. Membrane tubulation activity was also assessed for a N-terminally truncated variant of AtREM1.3 (AtREM1.3$^{\text{C-term}}$, J), a chimeric protein comprised of the AtREM1.3 N-terminal and the MtSYMREM1 C-terminal region (AtREM1.3$^{\text{N-term}}$::MtSYMREM1$^{\text{C-term}}$, K) and a dual-legume SYMREM1 chimera (LjSYMREM1$^{\text{N-term}}$::MtSYMREM1$^{\text{C-term}}$, L). Scale bars indicate 10 µm. All images are maximum projections of images obtained by confocal laser-scanning microscopy. The red signal indicates chlorophyll autofluorescence from chloroplasts.

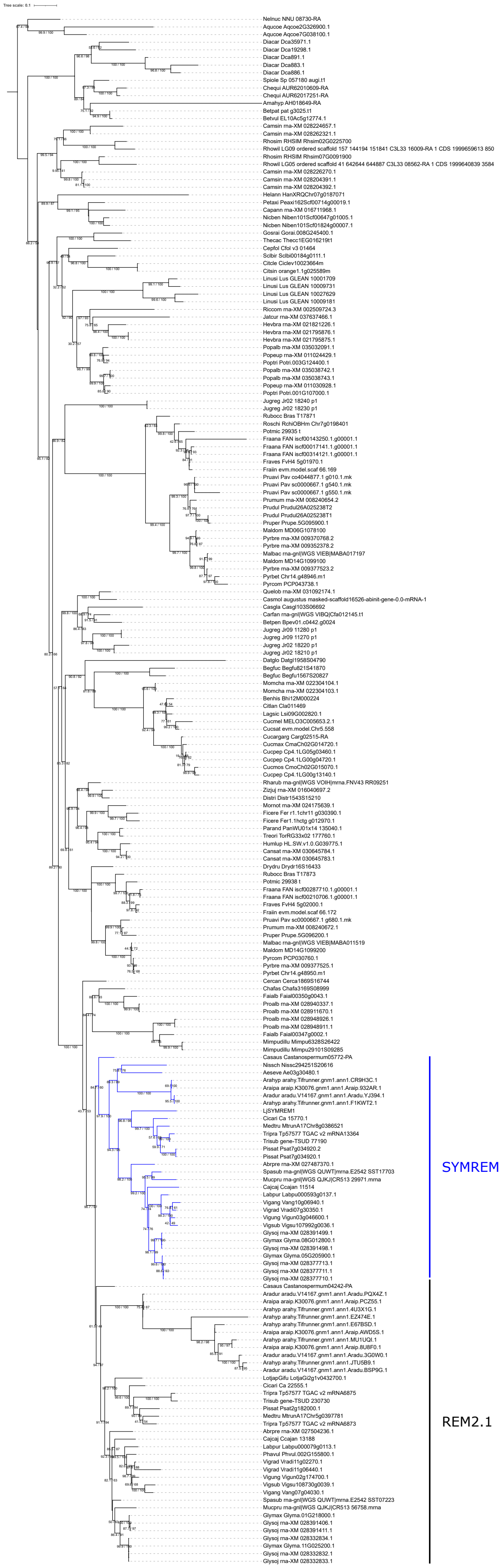


**Fig. S5. Maximum likelihood tree of SYMREM1 (co-)orthologs in Eudicots** (model: VT+R7; log-likelihood: -33187.1885). The tree is rooted on basal Eudicot species. Branches of direct orthologs of SYMREM1, deriving from the Papilionoideae duplication, are highlighted in blue. Branches supports are indicated as follow: sh-aLRT/UltraFastBootstraps. Species names are indicated by a six-letter prefix (see also Table S2).

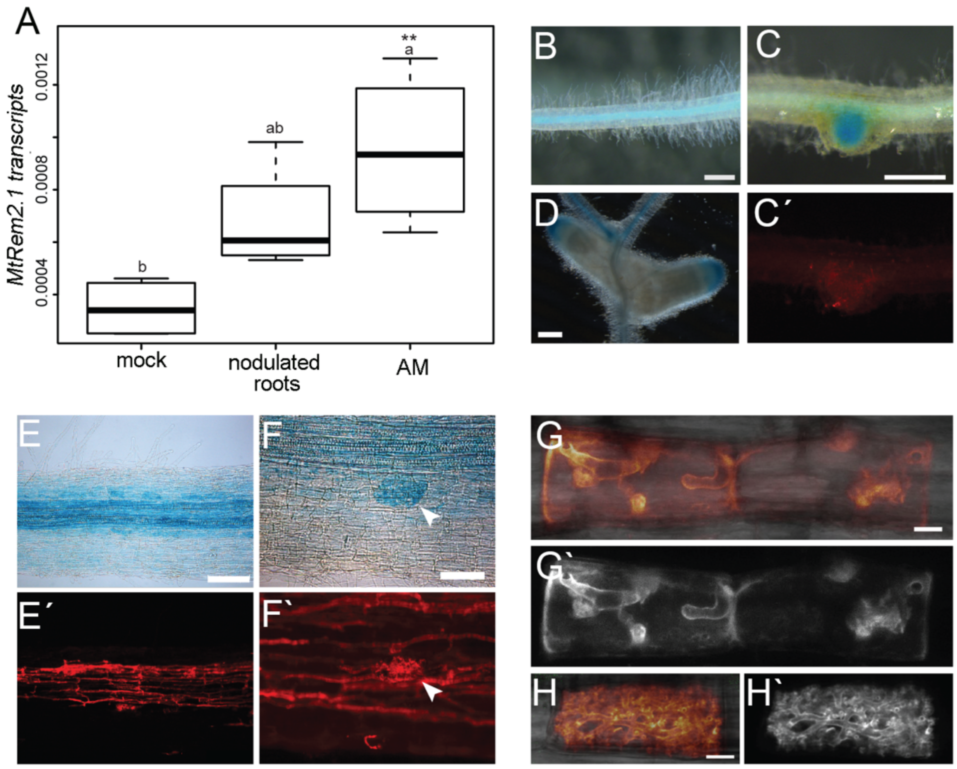


**Fig. S6. Expression and localization analysis of MtREM2.1.** (A) *MtREM2.1* transcripts were determined in whole roots (wild-type R108) 5 weeks after mock treatment (mock), after 4 weeks mock and 7 dpi with *S. meliloti* CFP (nodulated roots) or 5 weeks inoculated with *R. irregularis* (AM) (n=4). For statistical analysis a one-way ANOVA was performed followed by a TukeyHSD (significant differences indicated with small letters) and Dunnett test (significant differences indicated with an asterisk; p-value <0.05). (B-C') Analysis of *MtREM2.1* promoter activation using a *ProMtREM2.1::GUS* reporter in transformed roots of the R108 wild-type background. Weak GUS staining was detected in the vascular tissue under mock conditions (B). Upon inoculation with *S. meliloti*, GUS activity was found in uninfected nodule primordia 7-10 dpi (C and C') and was later restricted to the meristematic and distal zone II of 28 days old mature nodules (D). At 4-5 wpi with *R. irregularis*, fully colonized roots showed GUS activity in cortical cells and increased staining in arbuscule-containing cells (E, F). Fungal structures were stained with WGA-Alexa594 (E's, F'); arrow heads (in F and F') indicates an arbuscule-containing cell. (G-H') Localization pattern of mCherry-MtREM2.1 expressed under the control of the native *ProMtREM2.1* promoter in stable transgenic *M. truncatula* plants (wild-type R108) at 2wpi with *R. irregularis*. MtREM2.1 specifically labelled the symbiotic membrane surrounding intracellular hyphae with one or few branches (G and G') as well as extensively branched arbuscules (H and H'). (G-H') are maximum projection of confocal images, with G and H showing the overlay of the mCherry channel (false colored with LUT Glow) and bright field, (G' and H') show the mCherry channel. Scale bars=500µm (D, E and F), 100µm (G and H) and 10µm (I and J).
